# Sex-specific Proximal Tubular Cell differentiation pathways identified by single-nucleus RNA sequencing

**DOI:** 10.1101/2023.06.02.543031

**Authors:** Yueh-An Lu, Tanya Smith, Sumukh Deshpande, Chia-Te Liao, Bnar Talabani, Irina Grigorieva, Anna Mason, Robert Andrews, Timothy Bowen, Philip R. Taylor, Donald Fraser

**Affiliations:** Division of Infection and Immunity, School of Medicine, Cardiff University, Cardiff, United Kingdom; Wales Kidney Research Unit, Cardiff University, Cardiff, United Kingdom; Division of Nephrology, Kidney Research Center, Linkou Chang Gung Memorial Hospital, Linkou, Taiwan; Division of Nephrology, Department of Internal Medicine, Taipei Medical University-Shuang Ho Hospital, Taiwan; Division of Nephrology, Department of Internal Medicine, School of Medicine, College of Medicine, Taipei Medical University, Taiwan; TMU-Research Center of Urology and Kidney, Taipei Medical University, Taiwan; Dementia Research Institute, Cardiff University, Cardiff, United Kingdom

**Keywords:** cell biology and structure, chronic kidney disease, epithelial, kidney tubule, mRNA, proximal tubule, renal epithelial cell, renal fibrosis, renal tubular epithelial cells.

## Abstract

**Background:** Postpartum kidney growth is substantial but proliferation and differentiation pathways underpinning nephron elongation are not well defined. Here we performed sequential characterization of mouse kidney transcriptomics at the single cell level to address this.

**Methods:** Single nuclear RNA sequencing (snRNA-seq) was performed on kidney tissue from male and female mice at 1, 2, 4 and 12 weeks of age using the 10x Chromium platform.

**Results:** Unbiased clustering was performed on 68,775 nuclei from 16 animals. 31 discrete cellular clusters were seen, which were identified through comparison of their gene expression profiles to canonical markers of kidney cell populations. High levels of proliferation were evident at early time points in some cell types, especially tubular cells, but not in other cell types, for example podocytes. Proliferation was especially evident in Proximal Tubular Cells (PTCs) which are the most abundant cell type in the adult kidney. Uniquely when compared to other kidney cell types, PTCs demonstrated sex-specific expression profiles at late, but not early, time points. Mapping of PTC differentiation pathways using techniques including trajectory and RNA Velocity analyses delineated increasing PTC specialization and sex-specific phenotype specification.

**Conclusion:** Our single-cell transcriptomics data characterise cellular states observed during kidney growth. We have identified PTC differentiation pathways that lead to sex-specific tubular cell phenotypes.

## Introduction

The functional unit of the kidney is the nephron, which is dependent for its excretory and homeostatic functions on filtration of the plasma in the glomerulus, followed by selective reabsorption and secretion in tubular segments. The compartmentalized functions of nephron segments are reflected in the highly specialized phenotypes of the different cells contributing to overall nephron composition. Acquisition of these specialized phenotypes underlies nephron formation, a consequence of a series of reciprocal interactions between branching ureteric bud and pretubular aggregate occurring during development^1^.

Substantial postnatal growth in kidney size is evident, but underlying cellular processes are comparatively poorly categorized. Kidney growth occurs without increased nephron number in mammals, but nephron length is extended many-fold, and this is associated with extensive proliferation of Proximal Tubular Cells (PTC) and other tubular cell populations. PTC are quantitatively the greatest contributor to cellular composition of the adult kidney, and recent studies employing single nuclear RNA sequencing (snRNA-seq) have uncovered subsets of PTCs with discrete transcriptomic profiles^2, 3^. Profiles matching anatomically distinct PTC segments (S1, S2, S3) and intermediary or overlap profiles are evident. Rare PTC phenotypes, including proliferating and dedifferentiated cells, are seen in injured kidney where they may be key to failed repair processes and to progression of renal fibrosis^3, 4^. However, such rarer phenotype PTC are also present in normal adult kidney, where their function is less well understood and their low abundance renders them challenging to study.

Important differences exist between the sexes in excretory function, regulation of blood pressure and other homeostatic functions of the kidney^5^, and in risk and outcome of kidney diseases^6^. Sex differences in acid-base homeostasis have been linked to structural and functional differences in male and female proximal tubules^7^, and in a recent study employing a multiomic approach on microdissected mouse PTCs, differences between adult male and female mouse kidney were demonstrated^8^.

Here, we have studied transcriptional profiles at single cell resolution in kidneys from growing mice. We performed snRNA-seq on whole kidney from male and female mice at one, two, four and twelve weeks of age. Profound proliferation and increased abundance of differentiating cellular phenotypes has enabled study of PTC differentiation pathways. We delineate undifferentiated phenotypes that appear early in the post-proliferation trajectory of PTC differentiation and are common between males and females, and later mature phenotypes exhibiting significant sexual dimorphism associated with alterations in the cellular metabolic profile.

## Methods

### C57BL/6 mice

Experimental work was carried out using male and female C57BL/6 mice aged 1, 2, 4, and 12 weeks old from Charles River Laboratories. The 1, 2 and 4 week old mice were bred in-house by mating the C57BL/6 adult mice. Mice were housed with free access to chow and tap water on a 12 hour day/night cycle in a specific pathogen free environment. After mice were euthanized, chilled PBS (1x) prefusion via left ventricle was performed before kidney harvest. Body weight, kidney weight and kidney length were recorded. Kidneys from two male and two female healthy mice at all ages were processed for snRNA-seq. Hematoxylin and Eosin staining was performed on formalin-fixed, paraffin-embedded kidney sections. Experiments were performed in line with institutional and UK Home Office guidelines under the authority of an appropriate project license.

### Tissue collection and isolation of nuclei

For nuclear isolation for the snRNA-seq experiments, the 1 and 2 week old mouse kidneys were processed using a whole kidney, the 4 week old mouse kidneys were processed using half of a kidney and the 12 week old organs were processed using a quarter of a kidney from each mouse. The kidneys were minced into <2 mm pieces and transferred to a Dounce tissue grinder containing 2 ml of lysis buffer (Nuclei EZ Lysis buffer, Sigma NUC101) supplemented with protease inhibitor (Sigma 5892970001) and RNase inhibitors (Promega N2615 and Life Technologies AM2696)). Kidneys were homogenized and transferred into 50 ml tubes containing 2 ml of lysis buffer and incubated for 7 minutes on ice. The lysed cells were filtered through a 40-µm cell strainer and centrifuged at 500*g* for 5 minutes at 4°C. The supernatants were removed, the pellets were resuspended in 4 ml of lysis buffer, incubated for another 7 minutes on ice, then centrifuged again at 500*g* for 5 minutes at 4°C. The supernatants were removed, the pellets were resuspended in 4 ml of wash & resuspension buffer (1xPBS, 1.0% BSA, and RNase Inhibitor (Sigma 3335399001)) and filtered through a 20-µm cell strainer. Samples were then processed immediately using the 10x Genomics single-cell library preparation protocol.

### Library preparation and RNA sequencing

Library preparation was performed using Chromium Single Cell 3LJ Reagent Kits v3.1 (10x Genomics). cDNA quality was evaluated by fragment analysis (5200 Fragment Analyzer System, Agilent). RNA sequencing was carried out using the Illumina NovaSeq System.

### Data processing and core analysis of snRNA-seq dataset

The sequencing data were processed using the zUMIs pipeline (version 2.3.0)^9^. The pipeline was used to first discard reads with low-quality barcodes and UMIs, and then to map reads to the mouse reference assembly (Mus_musculus.GRCm38.95). The barcode-gene matrix generated by zUMIs was analyzed using the R package, Seurat (version 4.3.0)^10, 11^.

In Seurat, cells for individual samples were retained if they contained ≧ 200 genes and genes identified in ≧ 3 nuclei. Data from all 16 mouse kidneys were merged. Cells were filtered again to remove nuclei expressing ≦ 600 genes or ≧ 7500 genes, or with mitochondrial gene expression ≧ 7.5%. The feature counts were normalized with scale factor = 10,000. The top 2,000 variable genes were identified and scaled and the principal component analysis (PCA) result of the scaled data was obtained. The data was then processed using Harmony (version 0.1.1) for batch effect correction^12, 13^. *FindNeighbors* and *FindClusters* functions were applied based on previously corrected principal components (PCs) to identify clusters of nuclei. T-Distributed Stochastic Neighbor Embedding (t-SNE) plot was used to visualize clustering results. DoubletFinder (version 2.0.2) was then used to exclude potential doublets^14^. After doublet removal, the merged dataset was again normalized, scaled, processed with PCA and Harmony. The number of PCs included in the downstream analysis was determined by identifying the knee point of the elbow plot generated after running the *JackStraw* procedure. *FindNeighbors* and *FindClusters* were performed for clustering. Final clustering results were visualized using Uniform Manifold Approximation and Projection (UMAP).

The cell cycle analysis used *CellCycleScoring* to identify cells in the G2/M and S status. Cells with G2M.Score > 0.15 and G2M.Score > S.Score were assigned as G2M status. Cells with S.Score > 0.15 and S.Score > G2M.Score were assigned as S status. Cell with G2M Score <0.15 and S.Score < 0.15 were assigned as G1/G0 phase.

### Proximal tubular cell subclustering analysis

The PTC clusters were subset for further analysis. PTCs of 1, 2, 4 and 12 week old mouse kidneys were integrated using Seurat. The integrated dataset was then scaled and processed with PCA, *FindNeighbors and FindClusters*. A total of 30,396 PTCs were re-clustered and analyzed. A cluster containing 242 cells that expressed both PTC and endothelial marker genes and had higher genes per cell was removed as highly suspicious of residual doublets. Marker genes of the new PTC clusters were identified using differential gene expression (DGE) analysis. For DGE analysis, we used the *FindMarker* command. Significance was defined as a gene with an adjusted p value < 0.05, a ≧ 0.25 average log2-fold difference between the two groups of cells, and whose presence was detected in at least 10% of cells in either of the two populations. P-value adjustment was performed using Bonferroni correction based on the total number of genes in the dataset.

### Velocity and Pseudotime analysis

The RNA velocity of PTCs was calculated by velocyto.py and velocyto.R (version 0.6) using the spliced and unspliced RNA counts provided by 10X cellranger package^15^. The R package Monocle3 (version 1.3.0) was used for trajectory and pseudotime analysis^16^. The trajectory analysis of the re-clustered PTCs was performed using the *learn_graph* function. Pseudotime analysis was performed based on the RNA velocity result.

### KEGG pathway analysis

We conducted gene set enrichment analysis (GSEA) to understand pathways characteristic of the PTC clusters using the R package, WebGestaltR (version 0.4.3)^16^. We evaluated the pathway enrichment in the KEGG functional databases using the recommended False Discovery Rate (FDR) cutoff of 0.25 (https://www.gsea-msigdb.org/gsea/index.jsp).

### Immunofluorescence (IF) staining

Fixed mouse kidneys were processed for embedding in paraffin and cut into 5 μm sections for hematoxylin and eosin staining and immunofluorescence. Deparaffinized kidney sections were rehydrated in graded alcohols (100%, 96%, 70%, and 50%), and antigen retrieval was performed in citrate buffer in the autoclave at 120°C for 20 minutes. Sections were incubated with mouse-on-mouse block (Vector, MKB-2213-1) for 30 minutes and with 10% goat serum for an hour. Primary antibodies included anti-SLC5A2 (Biotechne, NBP1-92384), SLC13A3 (Biotechne, NBP1-82602), SLC5A10 (Biotechne, NBP2-13341) and Ki67 (Biotechne, NBP2-22112). The sections were incubated with primary antibodies overnight at 4 °C, then goat anti-mouse and goat anti-rabbit Alexa Fluor 488 or 594 conjugated antibodies (Invitrogen) were used as secondary antibodies. The sections were incubated with secondary antibodies for an hour at room temperature. Hoechst 33342 was used to stain the nuclei. Immunostained tissue slides were visualized and digitized using a confocal laser scanning microscope (LSM800, Carl Zeiss). Images were analyzed with the ZEN2012 software (Zeiss).

## Results

Kidney growth was evaluated in male and female mice. Macroscopic appearances of whole kidneys are shown (Fig 1A). Increases in body weight, kidney weight, kidney length and number of nuclei extracted from whole kidney were seen from 1 to 12 weeks in age and, by 12 weeks, males exhibited greater values for these parameters than females (Fig 1B). Histological appearances of female and male kidneys at each time point are depicted in Fig 1C.

**Fig. 1.**
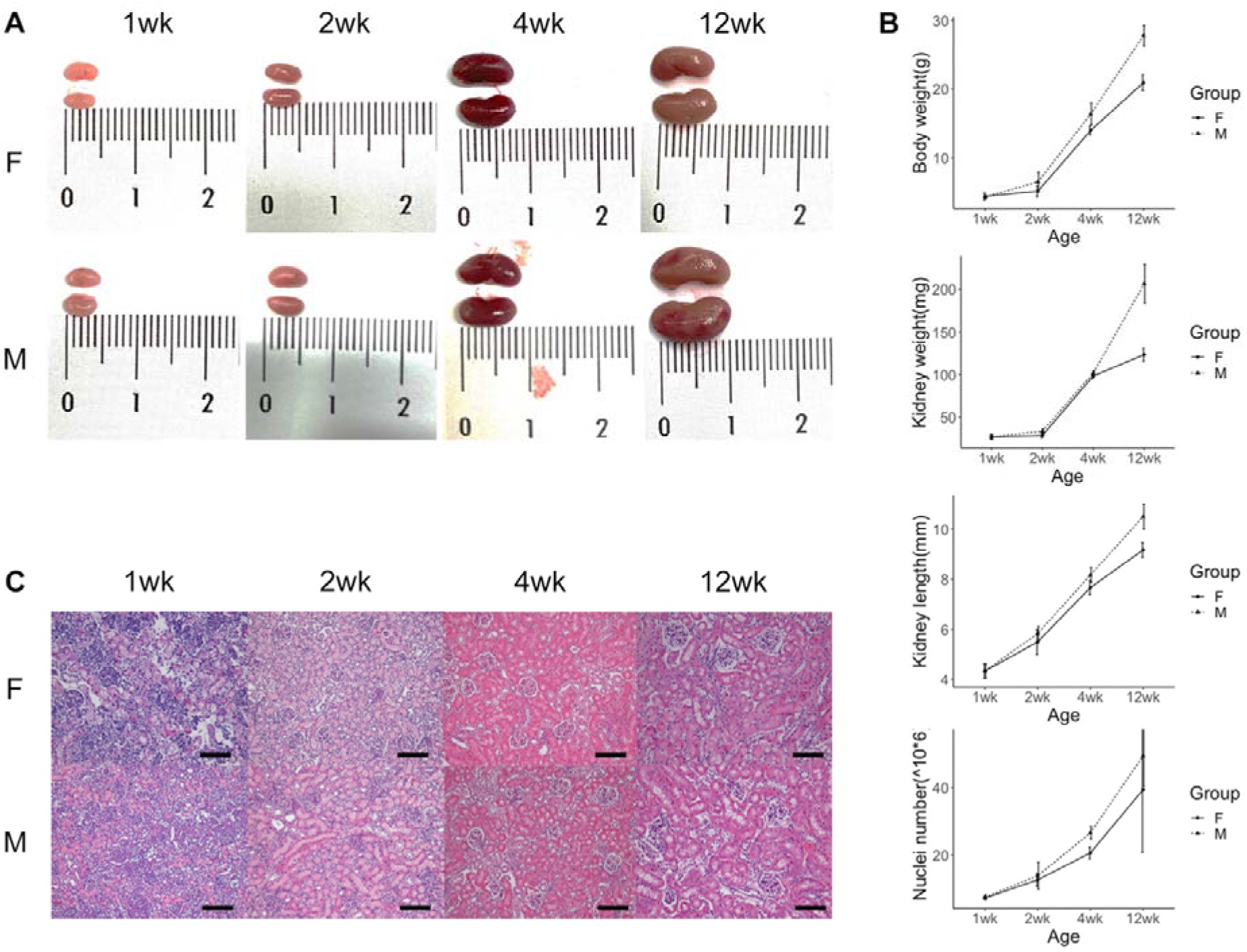
Appearances of female and male mouse kidneys at 1, 2, 4 and 12 weeks of age. (A) Kidneys freshly harvested from female (F) and male (M) mice at different ages. Ruler scale is shown in centimeters. The length and size of kidneys increased with age. (B) Line charts of body weight, kidney weight, kidney length and extracted number of nuclei from female and male mouse kidneys at different ages (n=3 of each group at each time point). Data is shown as mean +/- standard deviation. Adult male mice had higher body weight, higher kidney weight and kidney length and were comprised of more cells than adult female mice. (C) H&E stain of female and male kidneys at different ages (scale = 100μm). Tubular expansion was observed during growth.

SnRNA-seq was performed on kidney tissue from male and female mice at 1, 2, 4 and 12 weeks of age. Unbiased clustering was performed on 68,775 nuclei, revealing 31 separate clusters (Fig 2A, Supp Fig 1). Cluster identification was performed using canonical marker genes (Fig 2B). Kidneys of different ages varied in the proportional contribution to cellular number of each cluster (Fig 2A). Cell cycle status identified cells in G2/M and S phase versus those in G1 (Fig 2C). While tubular nephron segments exhibited abundant proliferation, proliferation of other cells was less evident. Alterations in overall cellular composition over time were evident, with some clusters contributing a greater proportion of total cell number at early time points, including proliferating cells of various lineages (Fig 2E, Supp Fig 3) as well as podocytes, mesangial cells, endothelial cells, juxtaglomerular cells and fibroblasts (Fig 2F, Supp Fig 3). In contrast, the proportion of total cell number that were tubular cells of various types increased throughout kidney growth (Fig 2G, Supp Fig 3).

**Fig.2.**
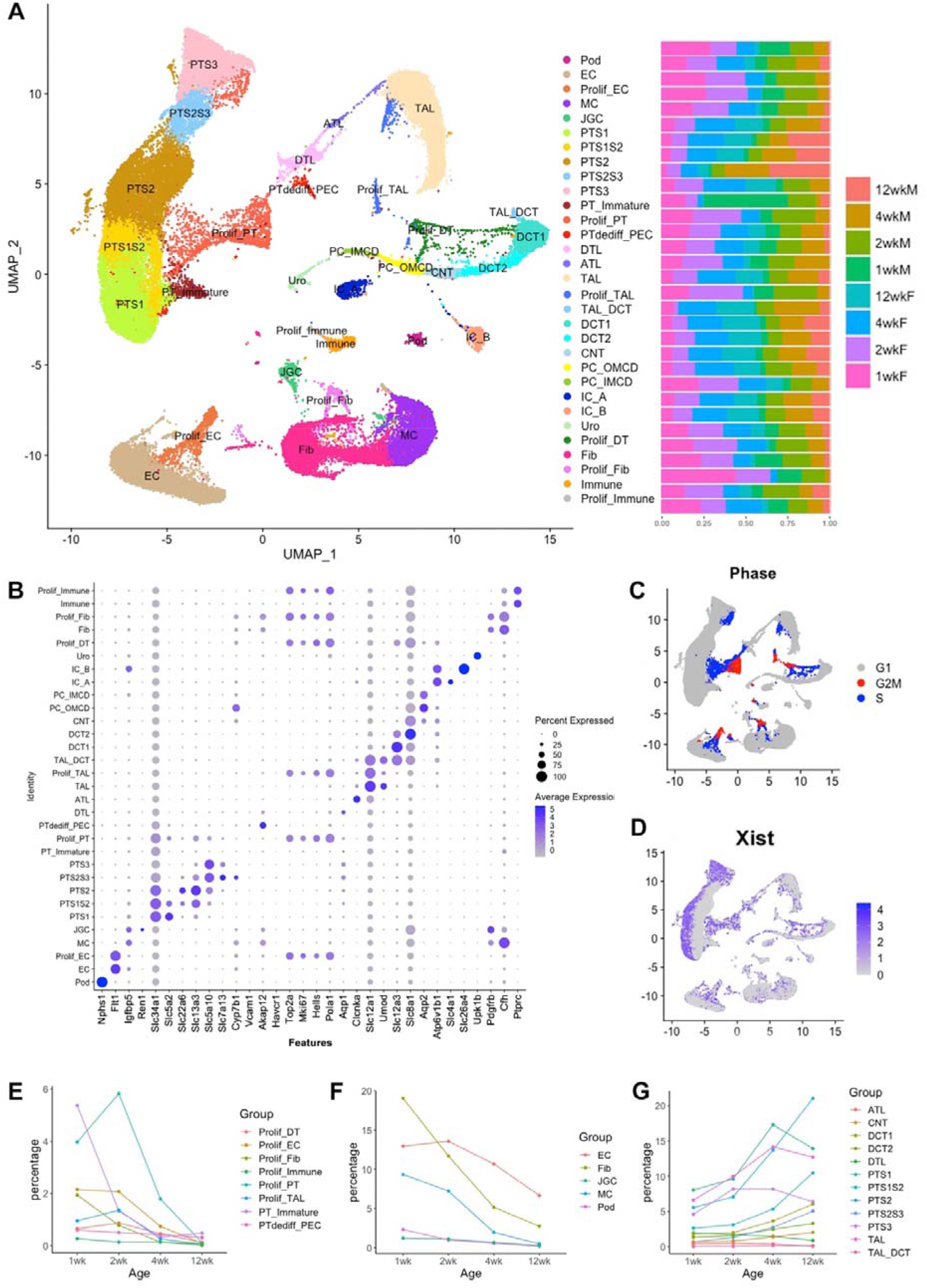
SnRNA-seqof 68,775 nuclei from female and male mouse kidneys at 1, 2, 4 and 12 weeks of age. (A) Results of cell clustering and cell type identification are shown using uniform manifold approximation and projection (UMAP) plot. Expected major types of kidney cells and their respective proliferative cell clusters were identified. (B) Dotplot shows the expression levels and the percentage of gene expression of the canonical genes in each cluster. (C) Results of cell cycle analysis. (D) Feature plot shows cells that express the Xist gene, which is specific for female cells. Regional expression of Xist is noted in PTCs but not in other cell types on the UMAP. (E/F/G) Evolutional changes of the percentage of each cell type at different ages. The percentage of proliferative cells, EC, fib, JGC, MC, Pod decreased with age, whereas the percentage of tubular cells increased with age. Pod, podocyte; EC, endothelial; Prolif_EC, proliferative endothelial cells; MC, mesangial cell; JGC, Juxtaglomerular cells; PT, proximal tubule; S1/S2/S3, segment 1/2/3 of proximal tubule; Prolif_PT, proliferative proximal tubule; PTdediff_PTC, dedifferentiated proximal tubule_parietal cell; DTL, descending thin limb; ATL, ascending thing limb; TAL, thick ascending limb; Prolif_TAL, proliferative thick ascending limb; TAL_DCT, thick ascending limb_ distal convoluted tubule; DCT1/DCT2, distal convoluted tubule1/2; CNT, connecting tubule; PC_OMCD, principal cell-outer medullary collecting duct; PC_IMCD, principal cell-inner medullary collecting duct; IC_A, intercalated cells, type A; IC_B, intercalated cells, type B; Fib, fibroblast; Prolif_Fib, proliferative fibroblast; Uro, urothelial cell.

The data demonstrated substantial numbers of proliferating PTCs at early time points, and increasingly dominant contribution to overall kidney cell number of PTCs at later time points (Fig 2E-G). Comparison of overall data further demonstrated differences between male and female animals in PTC clusters but not in clusters of other phenotypes (Supp Fig 2). Recent work from the Knepper laboratory has found important sexual dimorphism in microdissected proximal tubular segments using a multiomic approach^8^. Therefore, we next reclustered and analysed PTC. Before commencing this analysis, we sought to confirm the presence of proliferating cells across PTC segments. Ki67 was used as a marker of proliferating cells, while Slc5a2, Slc13a3 and Slc5a10 were used to identify PT segments S1, S2, and S3, respectively. Ki67-positive cells were found to be localized to S1, S2 and S3 PTC segments (Fig 3A-C).

**Fig. 3.**
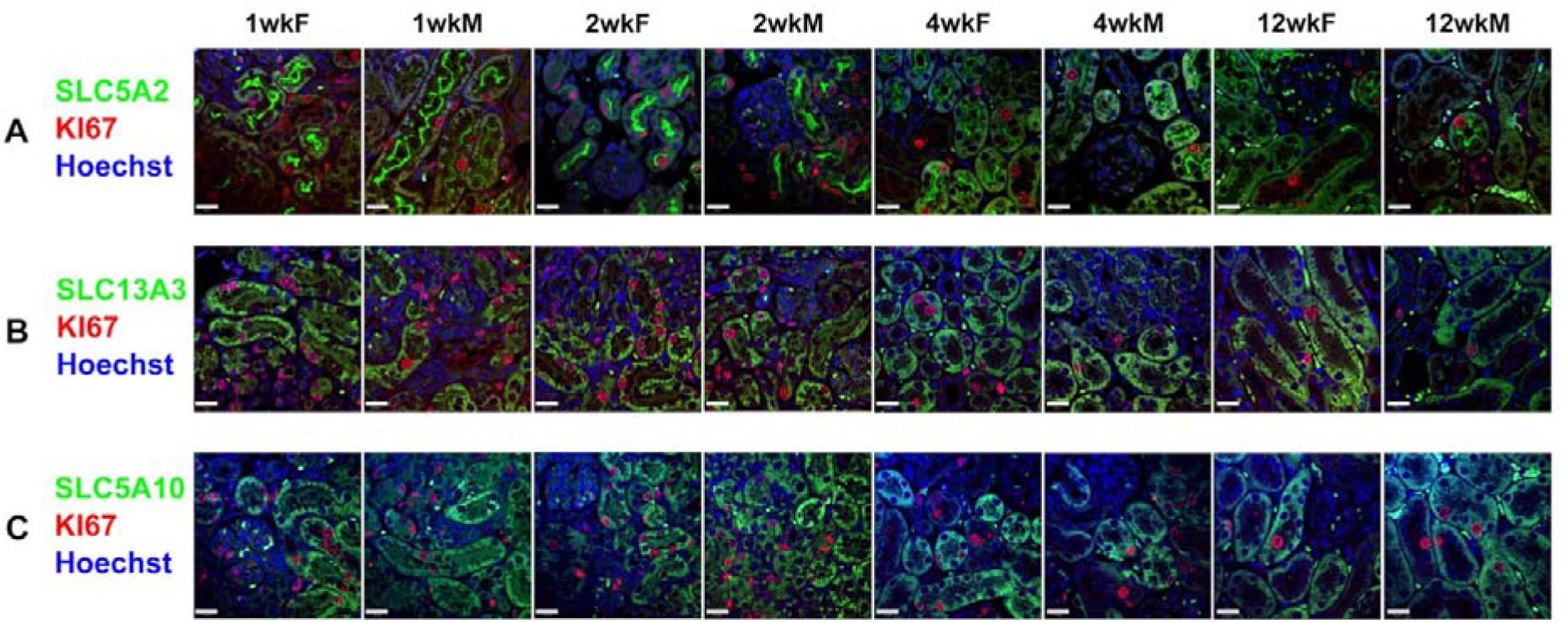
Validation of the snRNA-seq result using immunofluorescence stain of the proliferative cells in S1, S2 and S3 segments of the proximal tubule. Immunofluorescence stain shows the colocalization of the proliferative marker Ki67 and the PTC markers for S1 segment, SLC5A2 (A); S2 segment, SLC13A3 (B) and S3 segment, SLC5A10 (C) (scale = 20μm). Proliferating cells were identified in S1, S2 and S3 segments at 1, 2, 4, and 12 weeks-old male and female mice. The image is representative of 2 mice in each group. High proliferation was noted postnatally and reduced as the tissue matures in all three PT segments under microscope.

X-inactive specific transcript (Xist) is a long noncoding RNA that is expressed from the inactive X Chromosome in female cells, and functions as a central component of the X chromosome inactivation machinery that transcriptionally silences one of the two X chromosomes present in female cells^17^. Maternal microchimerism leads to the presence of a small proportion of maternal cells in the fetus, which persist in adult tissues and may have important developmental and functional consequences^18^. In this dataset, Xist was detected in 91.1% of cells from female animals and 0.3% of cells from male animals. Cells expressing Xist, whether from female mice or from male mice and of presumed maternal origin, demonstrated unequal distribution in the PTC clusters, but not in other nephron segments (Fig 2D).

Next, PTC reclustering analysis was performed (Fig 4A, Supp Fig 4). 15 PTC clusters were identified, of which 11 were comprised of Xist-positive and -negative cells, while 4 exclusively contained Xist-negative cells (Fig 4A, 4C, Supp Fig 5 and 6). Cluster annotation was performed using expression of canonical anchor genes (Fig 4B, Supp Fig 7 and 8). The transcriptional profile of the clusters identified cells of S1, S2, and S3 segment phenotype, together with “intermediate” phenotypes exhibiting S1/2 and S2/3 markers. Three separate proliferating clusters were observed, labelled “ProlifPT_1, 2, and 3”. Two undifferentiated clusters were seen, the first labelled “PT_Immature” and the second exhibiting overlap with a cluster that we have previously identified as comprising dedifferentiated PTC together with parietal epithelial cells^3^, here labelled “PTdediff_PEC”. Comparison of gene expression in male and female PTCs demonstrated comparatively few differentially expressed genes at 1 week and 2 week time points, but at 4 week and 12 week time points, increasing sexual dimorphism was evident (Fig 4D).

**Fig. 4.**
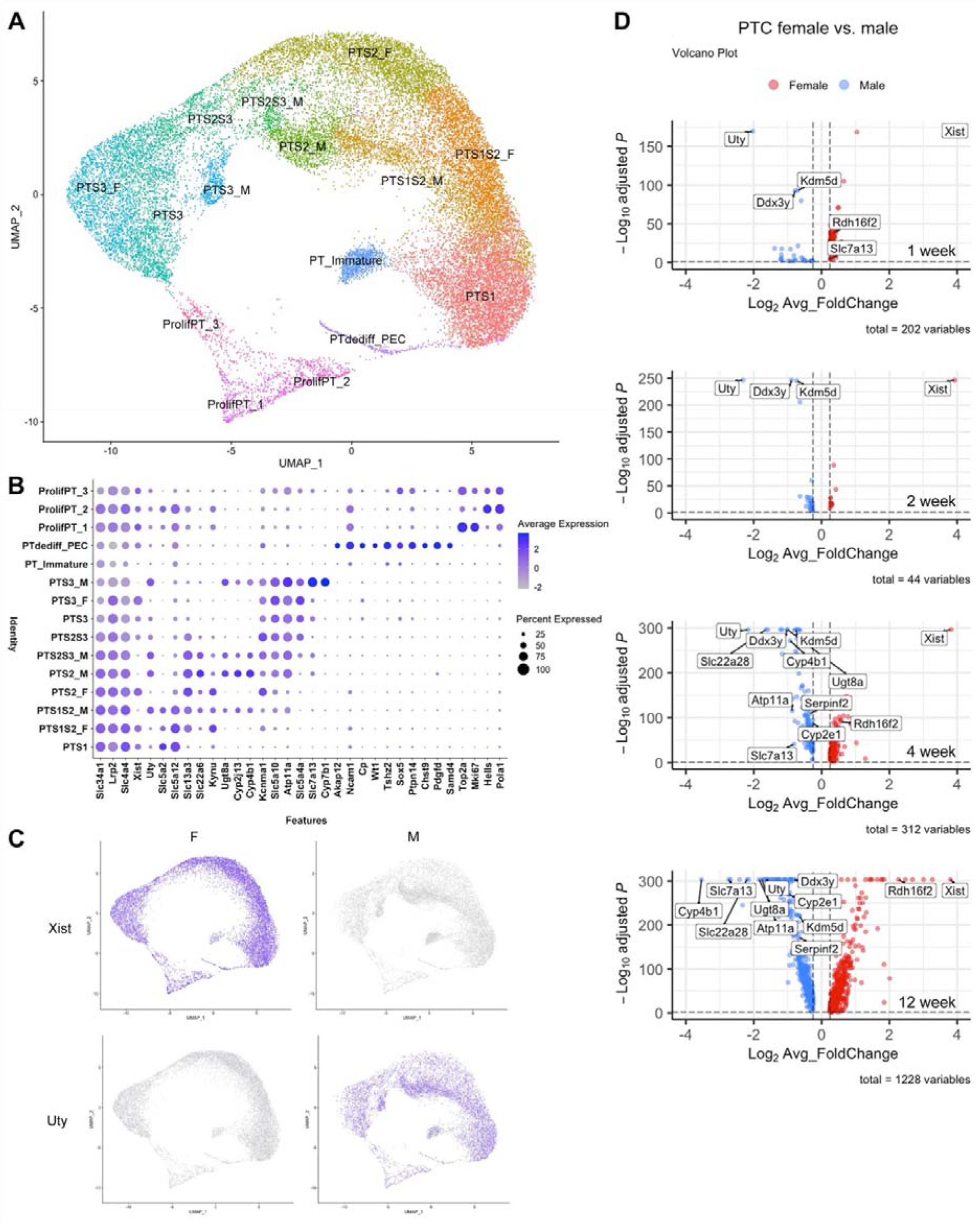
Analysis of PTCs of female and male mouse kidneys at 1, 2, 4 and 12 weeks of age. (A) Result of PTC re-clustering is shown using UMAP plot. (B) Dotplot shows the expression levels and the percentage of gene expression of the canonical genes in each of the PTC clusters. (C) Feature plot of the expression of Xist and Uty genes, which are specific for female and male cells, respectively. (D) Volcano plots show the results of differential expression gene (DEG) analysis of female and male PTCs at different ages. Twelve of the top 30 most differentially expressed genes are annotated, based on their comparatively high expression. These include Cyp2e1, Cyp4b1, Ddx3y, Kdm5d, Atp11a, Rdh16f2, Serpinf2, Slc22a28, Slc7a13, Ugt8a, Uty, and Xist.

Velocity analysis, which infers rate and direction of change in transcriptomic profile from systematic comparison of unspliced to spliced mRNA ratios^15^, was performed on the reclustered PTCs. Undifferentiated and proliferating clusters exhibited high levels of change in transcriptional profile, while the transcriptional profiles of mature PTC phenotypes appeared more stable (Fig 5A). Pseudotime analysis was used to determine PTC differentiation pathways.

**Fig. 5.**
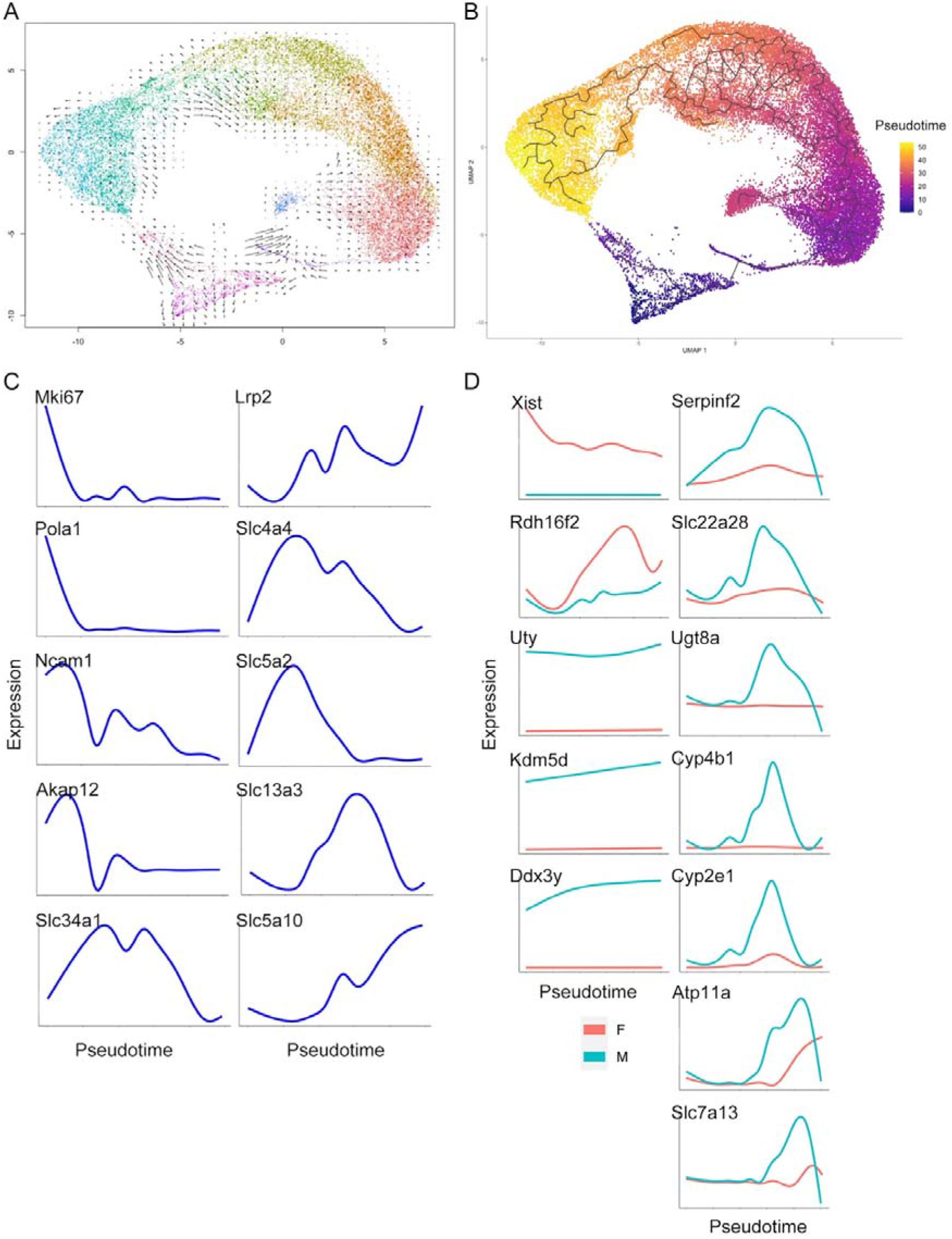
Velocity and Pseudotime analysis. (A) Velocity analysis shows that PTCs differentiation starts from proliferative PTCs and goes toward S1 and S3. (B) Result of pseudotime analysis. (C) Gene expression along pseudotime of marker genes of the PTC clusters. (D) Expression of genes selected from the DEG analysis, plotted along pseudotime in female and male mouse kidneys. Xist, Uty, Ddx3y and Kdm5d were persistently expressed throughout PTC differentiation whereas other genes demonstrate restricted expression patterns.

Ordering of cells along a latent (pseudotime) axis demonstrated a trajectory from proliferating through undifferentiated to mature phenotypes, the latter ordered anatomically from S1 to S3 (Fig 5B). Xist-specific clusters segregated late in the trajectory. Expression along the pseudotime demonstrated that following transient expression of proliferation-associated genes including *MKi67*, cells exhibited undifferentiated profiles followed by expression of markers of a fully differentiated phenotype (Fig 5C). The final, differentiated phenotypes were suggested by RNA Velocity analysis to be comparatively transcriptionally stable (Fig 5A). We next evaluated the 12 genes found to be abundantly expressed amongst those differentially expressed between male and female PTCs. 4 genes (*Xist, UTY, Kdm5d and Ddx3y*) were highly differentially expressed by male or female cells throughout, consistent with their localization on sex chromosomes and known roles in female/male cell specification^8, 19^. In comparison, genes originating on autosomes (*Rdh16f2, Serpinf2, Slc22a28, Ugt8a, Cyp4b1, Cyp2e1, Atp11a, Slc7a13*) were differentially expressed later in pseudotime (Fig 5D).

We next evaluated the high-level functions evident in the PTC clusters by KEGG analysis. Comparing 1 week and 12 week cells, 1 week cells exhibited enrichment for metabolic pathways, while 12 week cells were enriched in pathways including cell cycle regulation, adhesion, ECM interaction, and mineral absorption (Fig 6A). Male and female PTCs were similar at early time points but exhibited distinct transcriptional profiles at 12 weeks. KEGG analysis comparing 12-week male and female cells demonstrated that pathways comparatively upregulated in male cells included oxidative phosphorylation, arachidonic acid metabolism, metabolic pathways, and ribosome synthesis (Fig 6B). These data identify important differences in transcriptional profile, and in pathway activity between mature male and female PTCs.

**Fig. 6.**
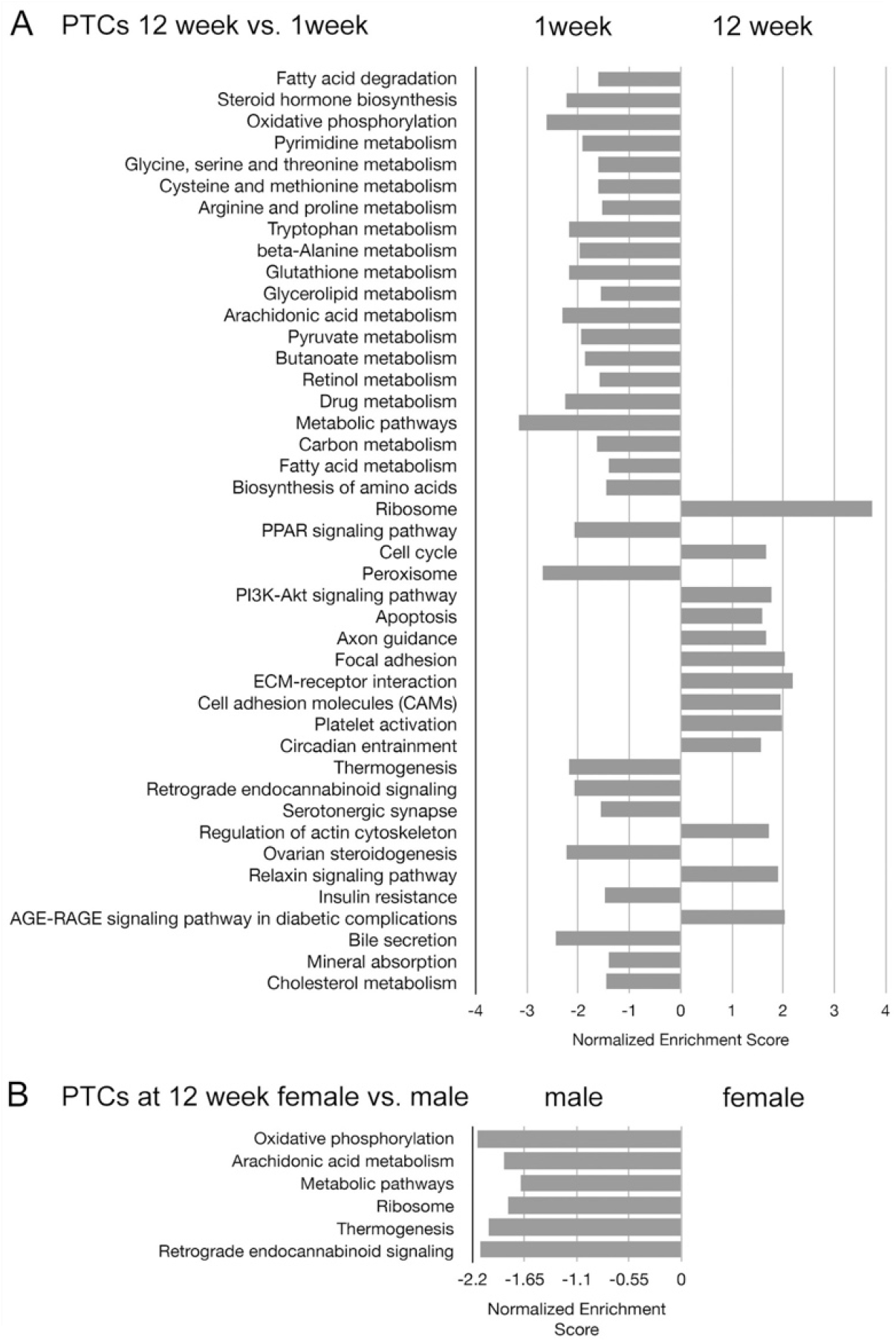
Pathway analysis of PTCs based on the KEGG database of (A) 12 week vs. 1 week and (B) female vs. male PTCs at 12 weeks old. Results show enriched metabolic pathway expression at 1 week old and enriched cell cycle and cellular community pathway expression at 12 weeks old. Pathways with a false discovery rate < 0.25 were listed. Male cells exhibited enriched energy metabolism and metabolic pathway expression when compared to female cells at 12 weeks old.

## Discussion

Here we have performed snRNA-seq on postnatal mouse kidney at time points from one to twelve weeks. In this time period, nephron number is fixed but kidney size, and individual nephron length, increase substantially. The data delineate the discrete cellular transcriptomic profiles present in the mouse kidney during postnatal growth. They further uncover important differences between mature male and female PTCs.

PTC and other differentiated kidney cell phenotypes arise during nephron formation. In the mammalian kidney, acquisition of the specialized phenotypes required for different nephron regions requires the reciprocal interactions of branching ureteric bud and pretubular aggregate^20^. Epithelialization of the pretubular aggregate forms the renal vesicle, from which the nephron then forms^21^. In humans all nephrons have been formed by 34-37 weeks’ gestation^22^. In mice, nephron formation has been observed up to postnatal day 4^20^.

Imaging studies and single cell RNA sequencing have shown that during nephrogenesis, progenitor cells are gradually recruited, and trajectory analysis demonstrates transcriptomic divergence as distinct regions of the nephron form^23^. Our data demonstrate important differences in postnatal cellular composition of the growing kidney, and suggest that postnatally, cellular differentiation continues at a different rate for different nephron regions and their respective cell populations. In support of this, previous studies have shown that while podocytes exhibit early commitment and terminal differentiation, tubular cells exhibit later and more complex differentiation pathways^24^. SnRNA-seq studies in adult mouse kidney have demonstrated unique profiles for tubular segments, together with presence of a quantitatively small proportion of cells exhibiting proliferative and less fully differentiated PTC phenotypes^2, 3, 25^. These minority phenotypes are seen in increased number following kidney injury, where additional pathology-associated phenotypes may be seen^3, 4, 26^. The rare nature of such cells in mature kidney outside of injury makes their profiling and determination of phenotype a challenge. The sequential profiling of growing kidney presented here, where proliferating and not yet fully differentiated phenotypes comprise a greater proportion of cell number, has uncovered the inter-relationships and development pathways of PTCs during kidney growth. These data map gene expression along a development pathway that proceeds from proliferation through intermediary phenotypes to cells with a mature, and anatomically restricted, expression profile.

Pathway analysis of PTCs based on the KEGG database identified comparatively high expression of metabolic pathways at early postnatal timepoints, whereas at later time points, PTCs demonstrated enrichment of pathways including those related to cellular adhesion and circadian entrainment. The data further demonstrate differences between cellular transcriptomic profiles in male and female kidney, that appear largely restricted to PTCs. Comparison of male and female PTC revealed relatively few differentially expressed genes at 1 and 2 week timepoints, while at 4 week and 12 week time points, increasing numbers of differentially expressed genes were found. Evaluation of sequential time points demonstrated sequential recruitment of genes that then remained differentially expressed in male and female PTCs at later time points.

Quantitative immunoblotting in adult rat kidney has previously demonstrated sex differences in renal transporters and electrolyte homeostasis^36^ and recent data from protocol transplant donor kidney biopsies identified increased metabolic activity of male PTC, supporting that this may also be found in human kidney^37^. In the current dataset, the genes identified as differentially expressed from 1 week notably included several genes located on sex chromosomes and with core roles in processes including X chromosome inactivation in female cells. 8 other genes, located on autosomes, were identified as abundantly expressed in kidney and exhibiting sexual dimorphism in PTCs at later time points (*Rdh16f2* in female PTCs, and *Serpinf2, Slc22a28, Ugt8a, Cyp4b1, Cyp2e1, Atp11a, Slc7a13* in male PTCs). These differences in PTC transcriptomic profile may underlie known differences in female versus male kidney responses to injury. For example, *Rdh16f2* expression, enriched in female PTCs at a late stage in the current dataset, is downregulated in male mice compared to female mice following ischemic injury^27^ and is linked to protective preconditioning in this context^28^.

Maternal michrochimerism may represent an additional source of complexity in the male versus female patterning of the proximal tubular component of the nephron. Maternal michrochimerism is the presence of a small proportion of cells of maternal origin in the progeny and is a result of vertical transmission of maternal cells to the fetus during mammalian pregnancy. The data presented here support the presence of maternal microchimerism in mouse kidney and suggest that in male animals, such maternal-origin cells retain transcriptionally female profiles in the proximal tubule. While the number of cells is quantitively small, data supports the importance of maternal microchimerism in patterning other organs and suggests that this warrants further study in kidney. For example, maternal michrochimerism has recently been found to promote brain development and homeostasis in mice^18^.

In summary, these data provide an atlas of transcriptomic data in the growing postnatal mouse kidney. They demonstrate that while postnatal cell proliferation is limited or absent for some cellular subtypes during mouse kidney growth, high levels of proliferation are present for others, including PTC. The results further highlight important sex differences in PTC differentiation pathways and differentiated PTC phenotype, which may have important implications for observed sex differences in outcome following acute kidney injury, experimentally and clinically.

## Data availability

## Acknowledgements

We would like to thank the staff of our animal facility for the care of the animals used in this study. P.R.T is funded by the Wellcome Trust Investigator Award (107964/Z/15/Z) and the UK Dementia Research Institute.

## Supplementary Data Table of Contents

