## Supplementary Data for "Sex-specific Proximal Tubular Cell differentiation pathways identified by single-nucleus RNA sequencing"

Supp Fig. 1 SnRNA-seq of 69775 nuclei from female and male mouse kidneys at 1, 2 ,4 and 12 weeks of age. Results of cell clustering and cell type identification are shown as uniform manifold approximation and projection (UMAP) plots at different ages.

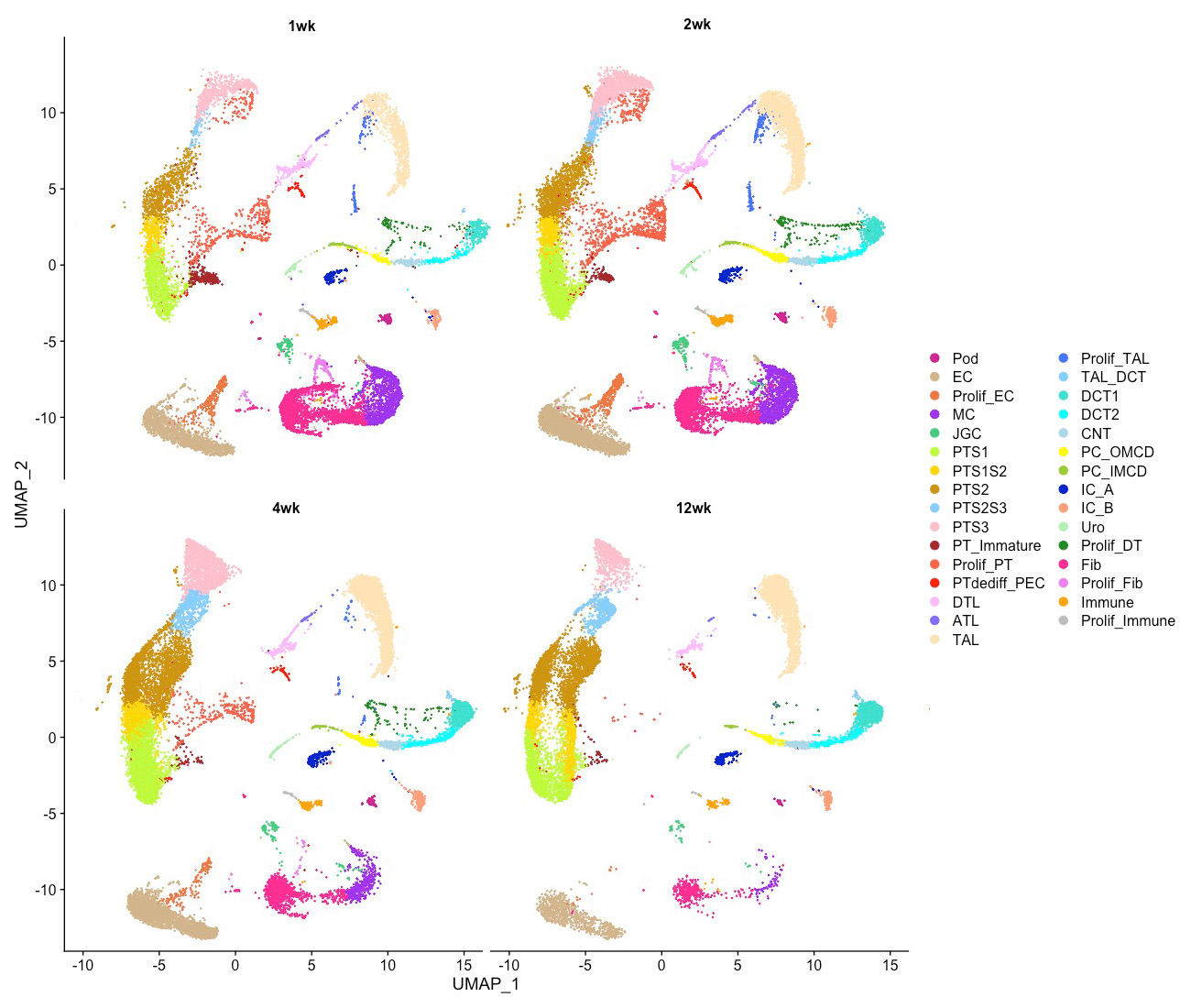

Supp Fig. 2 SnRNA-seq of 69775 nuclei from female and male mouse kidneys at 1, 2 ,4 and 12 weeks of age. Results of cell clustering and cell type identification are shown as female (F) and male (M) uniform manifold approximation and projection (UMAP) plots.

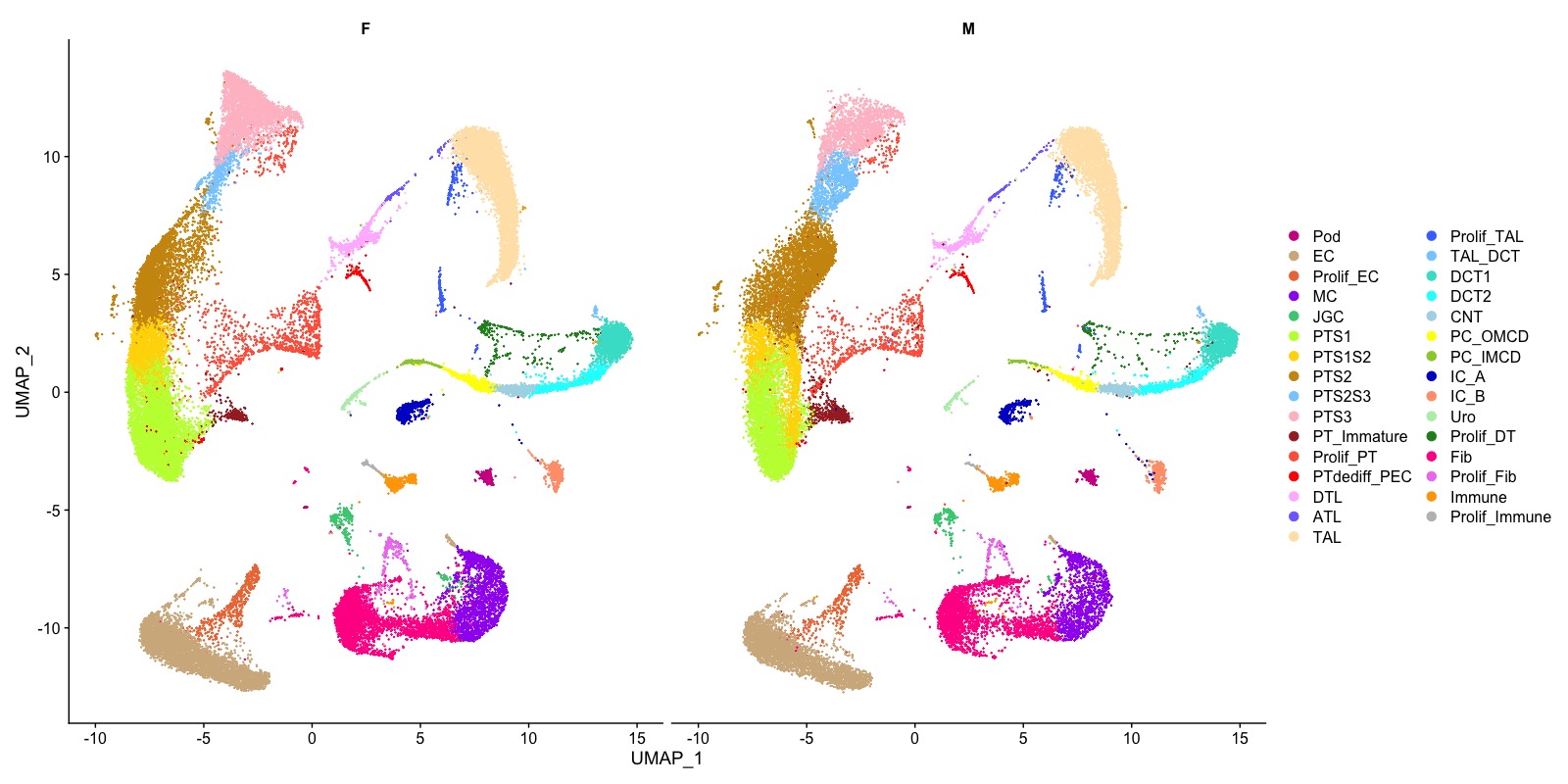

Supp Fig. 3 Line chart of the percentage of each cell type at different ages.

Pod, podocytes; EC, endothelial cells; Prolif_EC, proliferative endothelial cells; MC, mesangial cells; JGC, Juxtaglomerular cells; PT, proximal tubular cells; S1/S2/S3, segment 1/2/3 of proximal tubule; Prolif_PT, proliferative proximal tubular cells; PTdediff_PTC, dedifferentiated proximal tubular_parietal cells; DTL, descending thin limb cells; ATL, ascending thing limb cells; TAL, thick ascending limb cells; Prolif_TAL, proliferative thick ascending limb cells; TAL_DCT, thick ascending limb_ distal convoluted tubule cells; DCT1/DCT2, distal convoluted tubular ½ cells; CNT, connecting tubular cells; PC_OMCD, principal cell-outer medullary collecting duct cells; PC_IMCD, principal cell-inner medullary collecting duct cells; IC_A, intercalated cells, type A; IC_B, intercalated cells, type B; Fib, fibroblasts; Prolif_Fib, proliferative fibroblasts; Uro, urothelial cells.

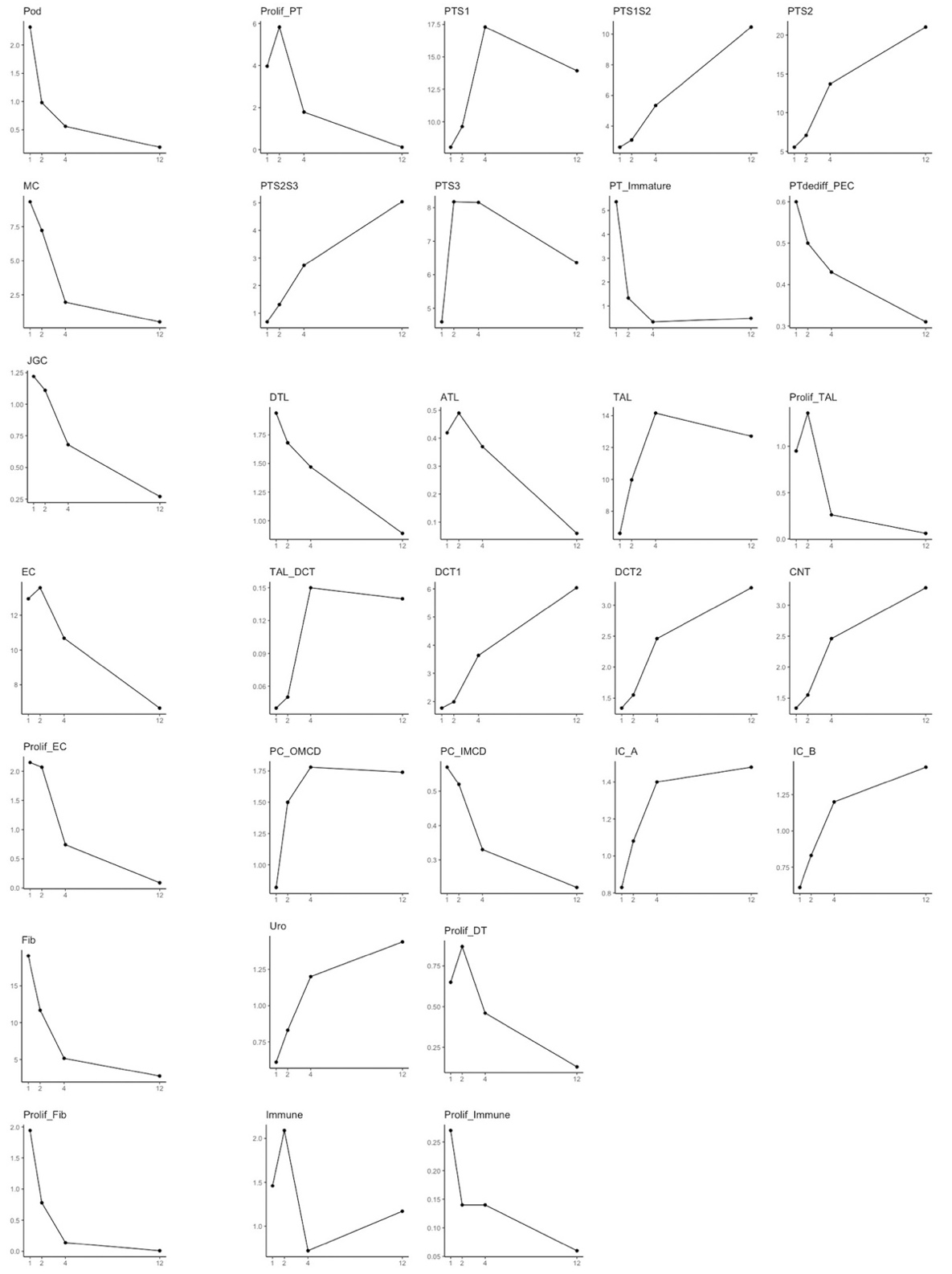

Supp Fig. 4 Result of PTC re-clustering. (A) UMAP plot shows the PTCs are discriminated into 25 clusters; (B) Results of re-cluster are compatible with the primary result of cell type identification; (C) Violin plots show the number of genes detected in each cluster. The proliferative clusters and cluster 25 contain greater numbers of genes per cell; (D, E) Feature plots of the expression of *Flt* and *Emcn* genes.

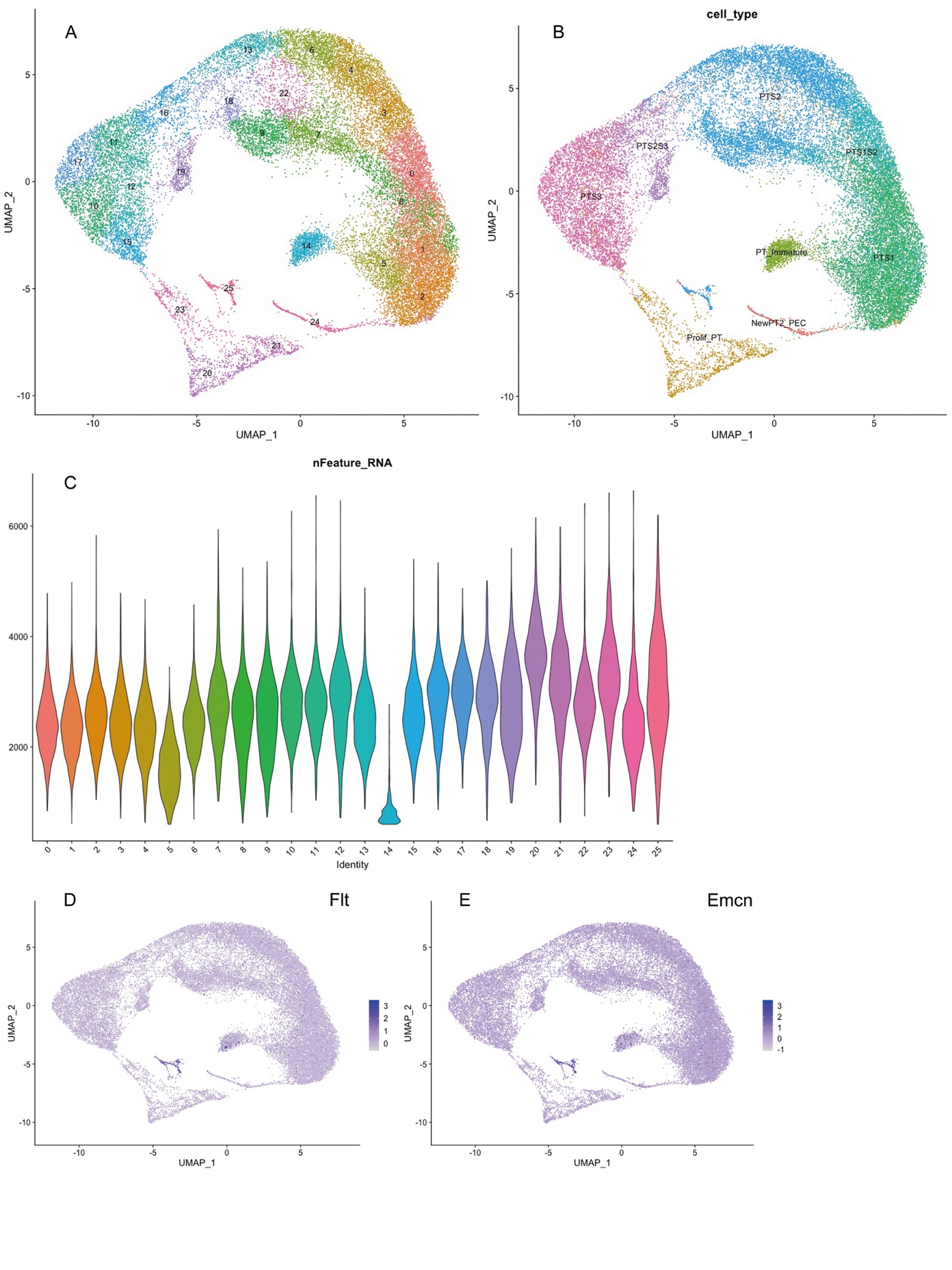

Supp Fig. 5 UMAP plots of re-clustered PTCs at different ages.

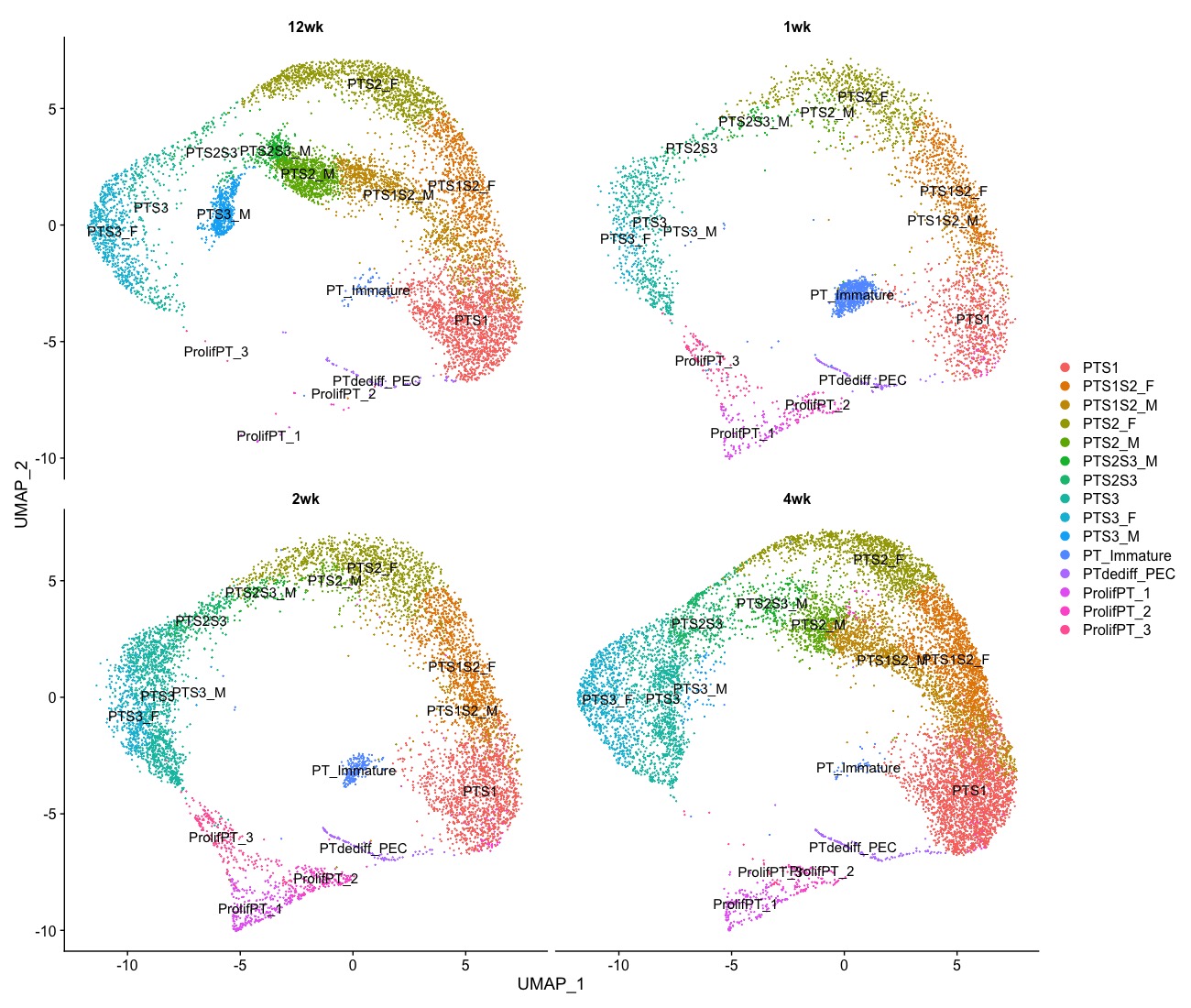

Supp Fig. 6 Female (F) and male (M) UMAP plots of re-clustered PTCs

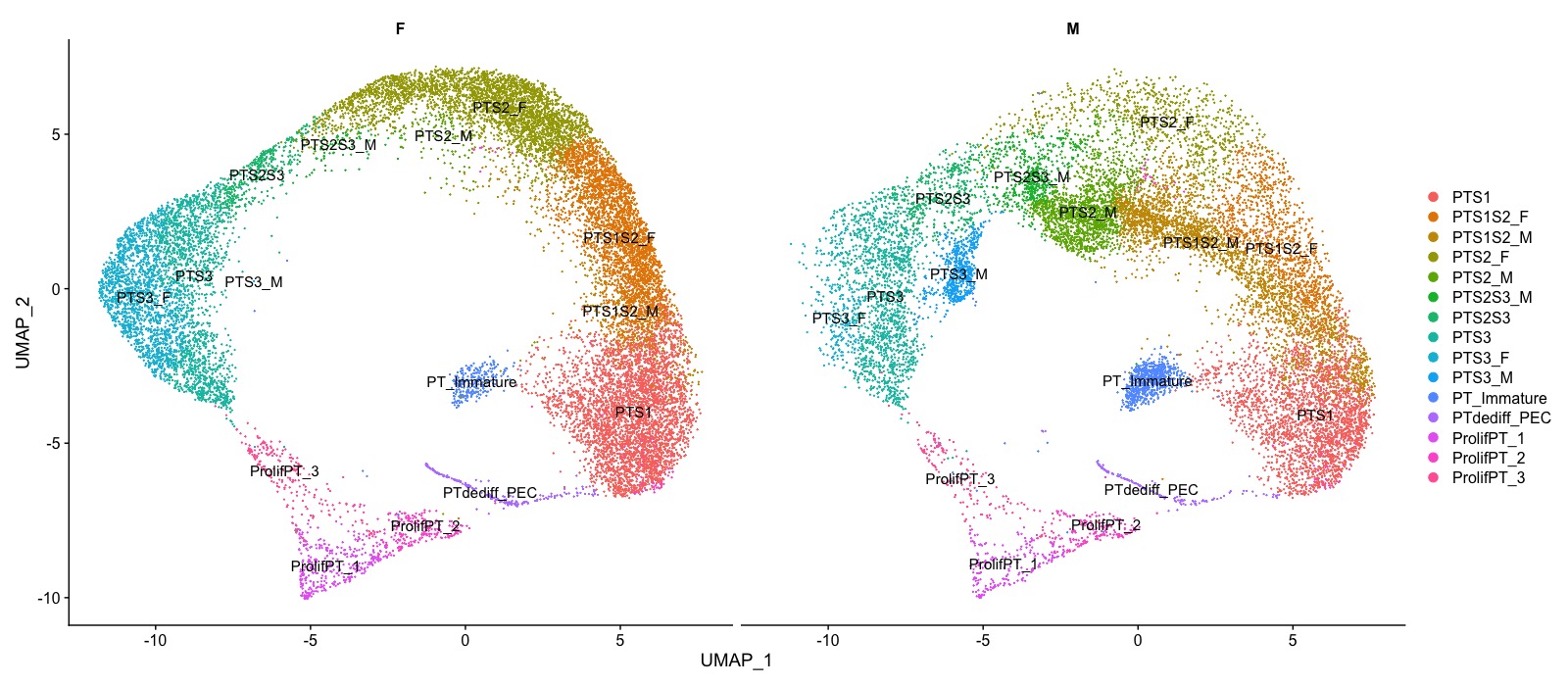

Supp Fig. 7 Dotplot showing the expression levels and the percentage of gene expression of PTC marker genes of female (F) and male (M) cells.

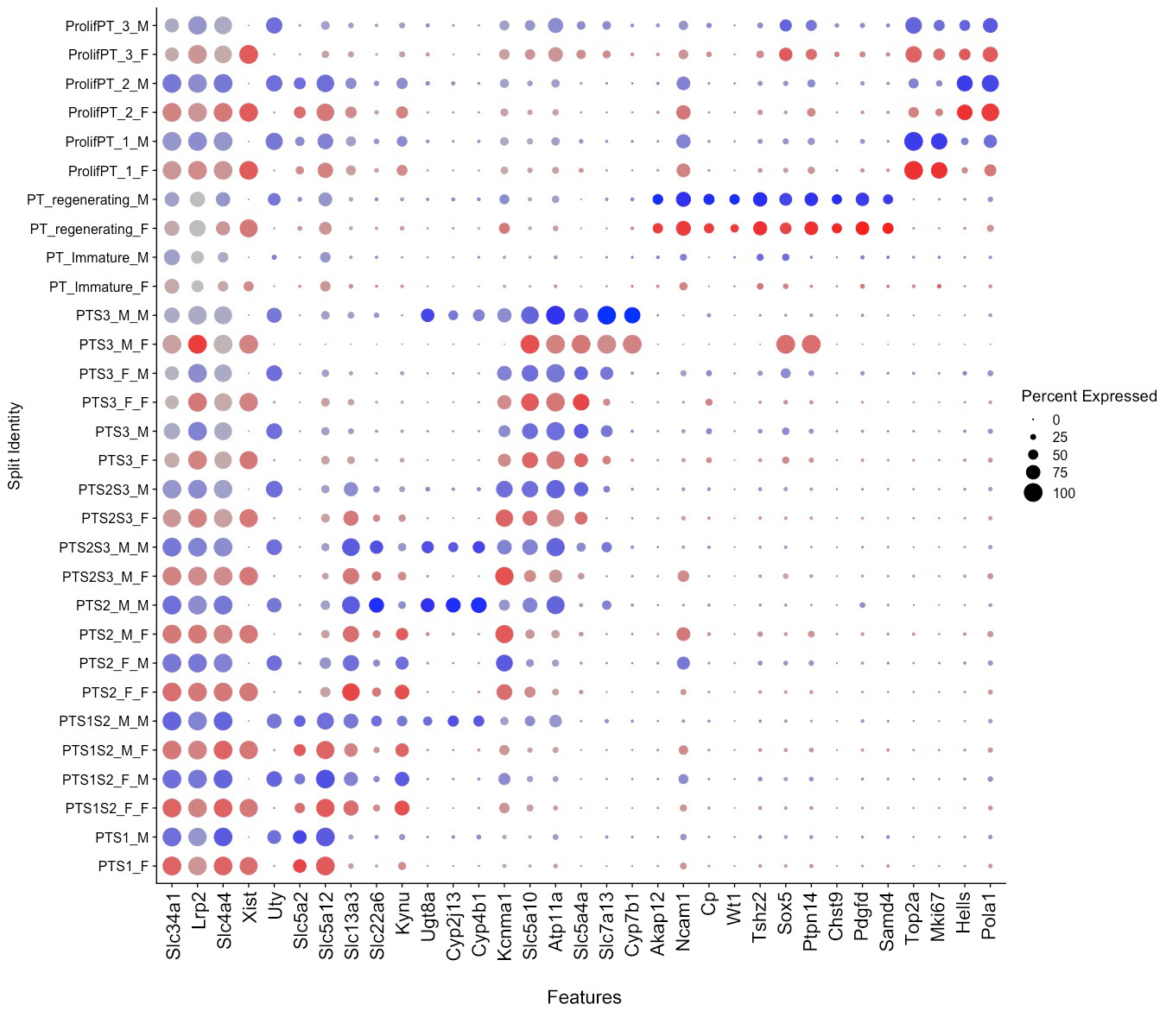

Supp Fig. 8 Dotplot showing the expression levels and the percentage of gene expression of PTC marker genes in all types of cells.

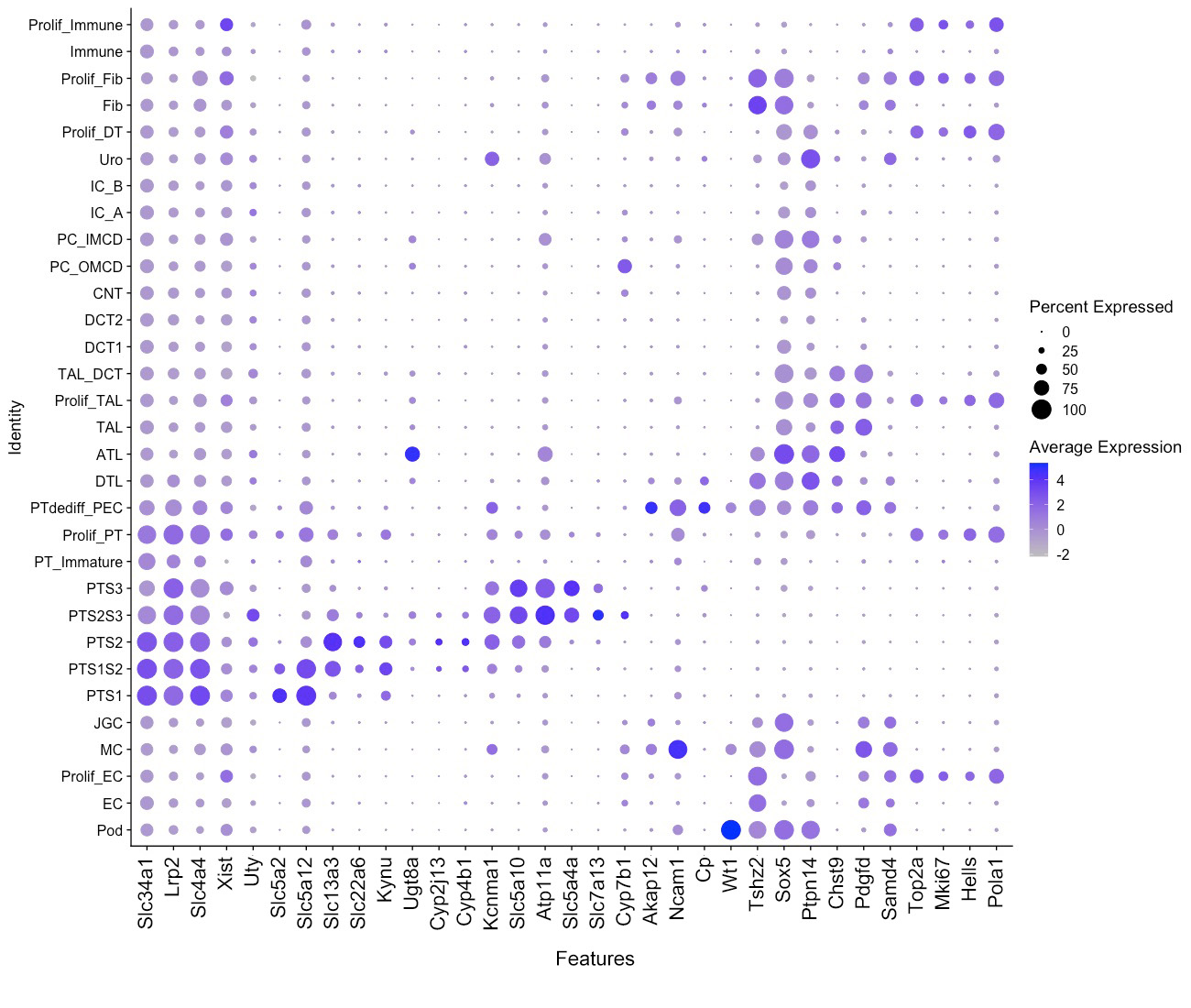
